# Receptor cavity-based screening reveals potential allosteric modulators of gonadotropin receptors in carp (*Cyprinus carpio*)

**DOI:** 10.1101/2023.10.12.562013

**Authors:** Ishwar Atre, Lian Hollander-Cohen, Hodaya Lankry, Berta Levavi-Sivan

**Author notes:** Corresponding Author: Berta Levavi-Sivan.

## Abstract

The gonadotropins follicle stimulating hormone (FSH) and luteinizing hormone (LH) are key regulators of sexual development and the reproductive cycle in vertebrates. Unlike most G protein-coupled receptors (GPCR), the FSHR and LHR have large extracellular domains containing multiple leucine-rich repeats, which leads to an elaborate mechanism of receptor activation via orthosteric sites that is difficult to manipulate synthetically. To bypass the orthosteric mechanism, in this study using carp as a model organism we identified allosteric sites capable of receptor activation on the transmembrane domain, which are spatially separated from the orthosteric sites. We have further generated pharmacophore hypothesis based on the structural motifs and exposed residues of these cavities. Using available online small compound libraries consisting of >70000 small molecules, we have thereon used receptor cavity-based hypothesis and other screening stages to identify potential modulators of the allosteric binding site on the carp FSHR and LHR *in-silico*. We then examined by *in vitro* transactivation assay the effect of four candidate compounds on FSHR and LHR, as compared to the activity of native ligands. Our results reveal both specific and dual effective allosteric modulators for FSHR and LHR, demonstrating the potential of our approach for efficient pharmacophore-based screening.

## Introduction

In vertebrates, the growth and activity of the gonads is regulated by two gonadotropins hormones (GTH): follicle-stimulating hormone (FSH) and the luteinizing hormone (LH). These hypophysiotropic hormones, belong to the glycoprotein family and play distinctive roles in reproduction. While FSH is responsible for gametogenesis and sustenance of ovarian follicles in females and sperm in males, LH is responsible for gamete maturation in both sexes and ovulation in females ^1, 2^. Both aquaculture and conservation depend on GTH hormones to regulate a species’ reproductive cycle for successful breeding and survival. However, in the absence of the natural ecosystems and environmental niches, hormonal secretion patterns affecting the reproductive cycle are perturbed, raising the need for synthetic manipulations of hormonal activity, by artificial molecular tools.

LH and FSH are heterodimers composed of a common glycoprotein α-subunit non-covalently attached to a unique β-subunit ^3^. These α-β complexes bind to gonadotropin receptors (GTHRs), which belong to the G protein-coupled receptor (GPCR) super family, to further influence the progression of gonadal development and sexual maturation. In mammals, knockout or dysfunctionality of GTHRs cause infertility and other complications related to vertebrate reproductive cycles, such as Hypogonadotropic hypogonadism and premature or delayed puberty, infertility, etc. ^4^. A majority of these dysfunctions can be overcome by synthetic or pharmacological stimulation of the receptor in question. In fish, loss of function of FSHR causes masculinization and suppression of ovarian development in female medaka, whereas LHR knockouts were observed to stop ovulation ^5, 6^. Recent studies leveraging advances in in silico methods and the availability of whole-crystal structures of GTHRs ^7, 8^ have greatly increased our understanding of GTH-GTHR binding mechanism and activation. In general, these receptors are seen to show activity when the cognate stimulant/ligand binds to their biologically active binding region called orthosteric binding site. Although the mammalian GTHR homologs in mammals display high specificity to their cognate hormones, in case of fish, both FSHR and LHR exhibit species-specific variability in ligand-binding promiscuity/specificity, which complicates the use of native ligands for receptor regulation. For example, in carp, FSHR is seen to be activated in response to both cFSH and cLH ^9^, whereas in tilapia, both GTHRs bind specifically to their cognate receptors ^10^. Simultaneously, in sturgeon ^11^ and medaka ^12^, both FSH and LH can activate each the others cognate receptor in addition to their own. Despite being the center of GPCR-targeted drug development, receptor modulation via orthosteric sites often lacks binding specificity, efficiency, and efficacy. Moreover, due to the large size of GTHR orthosteric sites, small compounds are unable to bind them and thereby change their conformation, impeding the use of low molecular weight drugs for pharmacological regulation of these receptors.

As opposed to most class A GPCRs that possess an orthosteric binding domain in the hydrophobic pocket created by the extracellular domain (ECD), the transmembrane domain (TMD), the connecting extracellular loops (ECL), GTHRs belong to a subfamily of glycoprotein receptors is with an ECD that is nearly as large as the TMD. This domain contains a series of leucine-rich repeat (LRR) domains that act as the orthosteric binding site, independently of the TMD. Furthermore, the hinge region which connects the ECD to TMD acts as a intramolecular modulator located between the ECD and TMD is thought to be essential for specificity in GTH-GTHR interaction. ^13^. Upon binding of the cognate hormone to the LRR region, this part moves to interact with the hinge region, which then manipulates the TMD and ICD, leading to receptor activation and downstream signal transduction ^7, 8^. Besides the orthosteric binding sites, several allosteric binding sites have been localized on GPCRs. Allosteric sites may lie within orthosteric binding pockets, overlap with them, or be topographically distinct. They can be located on the ECD, inside and/or outside the TMD and on the extracellular loops (ECL) or intracellular loops (ICL) of the receptor. Furthermore, these sites can accommodate agonistic, antagonistic, positive, negative as well as silent modulators whilst enabling the manipulation of multiple signaling cascades simultaneously. The allosteric sites on GTHRs are located in hydrophobic cavities formed by the TMD and ECL (Fig. 1) and are not biologically active or accessible for the native GTH, but can poses as common potential targets for small compounds that are highly effective reproductive endocrine modulators. While Allosteric sites in most GPCRs often partially overlap the orthosteric binding pocket in GTHR they are completely separated due to their unique ECD.

**Figure. 1:**
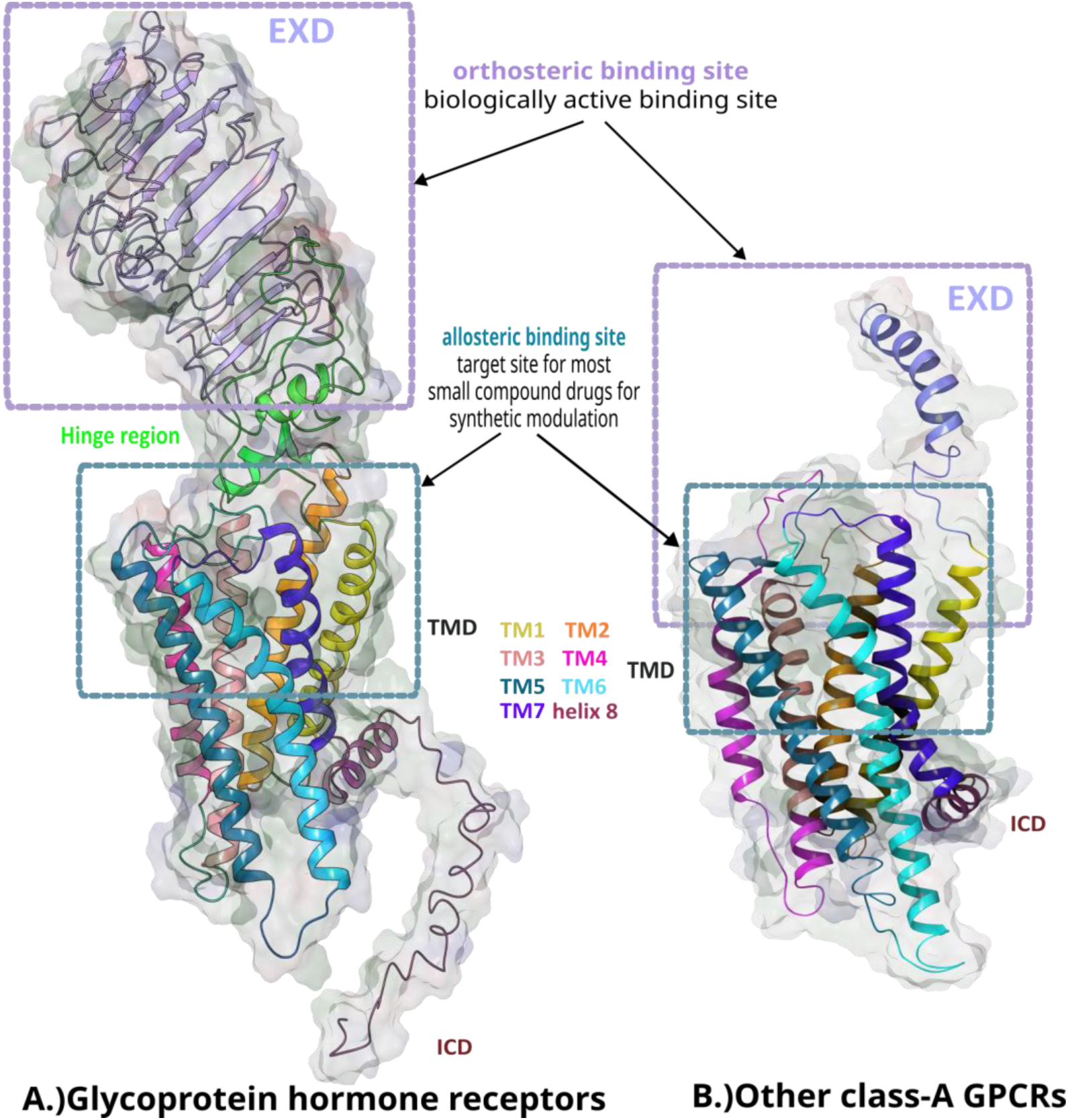
Orthosteric and allosteric sites in glycoprotein receptors vs other class A GPCRs. Ribbon diagrams show a glycoprotein receptor (A.) where the orthosteric site is situated on a large ECD that is seperated from the TMD and does not overlap the allosteric site, as observed in other class A GPCRs (B.).

A number of novel allosteric modulators have been reported for GTHRs over the past decade. However, most of them target human FSHR (hFSHR) and may not be as efficient or effective on GTHR homologs in other vertebrates like fish, which exhibit much more complex hormone-receptor expression, interaction, and specificity. Based on their effect on signal transduction and efficacy, the Allosteric modulators can be divided into four classes: agonists or antagonists that can directly modulate receptor activity and induce signal transduction without the involvement of additional ligands; positive (PAMs) and negative allosteric modulators (NAMs) that can potentiate or reduce native ligand-mediated response and thereby play a supportive role; and neutral allosteric ligands (NALs), which do not affect receptor activity after binding ^14–17^. A fifth category of modulators (Biased Allosteric Modulators), which has recently emerged, is defined by the signaling pathway-specific effects of the modulators on binding to designated receptors ^18^. Thiazolidines, which were the first GTHR-specific allosteric modulators to be discovered, bind exclusively to FSHR, showing no affinity to other glycoproteins such as LHR or thyrotropin receptor ^19, 20^. Nevertheless, some thiazolidine analogs may exhibit biased signaling and mobilize either Gα_s_, Gα_i_ or both. Similarly, recent *in vivo* studies have identified TP22 ^21^ and Org43553 ^22^ both of which are thieno[2,3-d]pyrimidine compounds as effective allosteric agonists of LHRs but might potentially influence different signal cascades downstream.

To date, most published studies focus mainly on FSH modulators, with most reported compounds being PAMs and NAMs. However, the screening process for allosteric modulators is lengthy and costly, which calls for the necessity of novel *in-silico* tools to increase the hit rate of the screening. However small compounds that modulate allosteric sites must overcome shallow binding pockets, low binding affinity, desensitization or mutational resistance, dissatisfactory ADME (absorption, distribution, metabolism, and excretion) values and the possibility of multiple site affinity ^23^. These limitations have called for the development of new tools to increase the hit rate and efficiency of screening. With the development in the field of *in-silico* tools, the candidate selection process and hit rates for these compounds have improved significantly. In this study, we have generated site-specific pharmacophore hypothesis based on the allosteric cavity at the site of interest. Combined with multiple *in-silico* screening methods, this approach enabled us to identify effective small compounds with high potential to act as agonists for carp GTHRs. Our *in vitro* results have confirmed that these compounds are independent modulators of GTHRs with high receptor specificity.

## Materials and Methods

### Homology modeling

3D homology models for carp cFSHR and cLHR were generated using hLHCGR homologs (PDB: 7FIG; 7FIH; 7FII; 7FIJ) as a template for both inactive and active states using the I-TASSER server *in silico* (Zhang 2009; Roy et al. 2012). The top models were selected based on C-score, structural stability, and structural similarity with the gonadotropin receptors. The protein models were further rendered and prepared using Maestro tool in Schrodinger software (Maestro, Schrödinger, LLC, New York, NY, 2021.).

### Pharmacophore model hypothesis, ligand screening and docking

Potential binding sites in the cFSHR and cLHR TMDs were detected using SiteMap module (Schrödinger, LLC, New York, NY, 2021) and selected based on their position in the TMD cavity. We then used these data to generate receptor cavity-based pharmacophore hypothesis. GPCR library (version 24 May 2020; https://enamine.net/compound-libraries/targeted-libraries/gpcr-library) (∼54000 compounds)) and allosteric GPCR library (version 28 February 2019; https://enamine.net/compound-libraries/targeted-libraries/gpcr-library/allosteric-gpcr-library) (∼14400 compounds) were downloaded from Enamine website (Enamine Ltd). The compound libraries were converted to phase databases and then screened using the Phase module of Schrodinger software, based on the generated receptor cavity-based and ligand-based hypothesis^24^. The ECD domain, excluding most of the hinge region, was cleaved off, and ligands were docked only onto the transmembrane allosteric binding pocket on carp GTHR using GLIDE module of Schrodinger software ^25^. To further screen the candidates, we used QikProp (Schrödinger, LLC, New York, NY, 2021) to predict ADME properties. Docked ligands were then further screened using g-score, docking score and e-model score, resulting in the selection of 20 small compounds for both cFSHR and cLHR. Four small compounds were eventually selected for *in vitro* studies based on predicted structural alignment to the receptor cavity. The interactions between the selected ligands and receptors were then analyzed *in silico* and compared to activation by orthosteric ligands.

### LUC Transactivation assay

Transient transfection, cell procedures and stimulations were generally performed as described previously ^10, 26, 27^. Briefly, COS-7 cells were grown in DMEM supplemented with 10% FBS, 1% glutamine, 100 U/ml penicillin, and 100 mg/ml streptomycin (Biological Industries, Israel) under 5% CO_2_ until confluent. The selected compounds (Z2242908028 (here on “8028”), Z1456504681(here on “4681”), Z2242909045(here on “9045”), Z1456630801(here on “0801”)) (Supplementary Spreadsheet. 1) were purchased from Enamine (Enamine Ltd., Kyiv, Ukraine). Co-transfection of the receptors (at 3 µg/plate) and cAMP response element-luciferase (CRE-Luc) reporter plasmid delivery (3 µg/plate for cFSHr and 0.3 µg/plate for cLHr) was carried out with TransIT-X2® System (Mirus). The cells were serum-starved for 16 h, stimulated with the various stimulants (initial concentration of 2 µg/ml diluted continually 1:3) for 6 h, and then harvested and analyzed. Recombinant carp LH (Aizen et al., 2012) and FSH (Hollander-Cohen et al., 2018) were used as positive controls. Lysates prepared from the harvested cells were assayed for luciferase activity, as described previously ^27^. Experiments were repeated at least three times from independent transfections, and each was performed in triplicate. Non-transfected COS& cells were used as negative control to ascertain the activity of the ligands is receptor specific.

### Statistical analysis

EC_50_ values were calculated from concentration-response curves by means of computerized nonlinear curve fitting (log(agonist) vs. response (three parameter)) using GraphPad PRISM 9 (version 9.5.0). The potency ratio was calculated as the Log(Relative Potency)= Log(EC50 of the native compound) – Log(EC50 of the novel compound) ^28^.

## Results & Discussion

To generate homology models for carp FSHR and LHR we used available crystal structures of human homologs (PDB ID: 7FIH & 8I2H, for cLHR and cFSHR, respectively) ^7^ ^29^, as the carp homologs displayed high similarity and features characteristic of human GTHRs. These glycoprotein receptors belong to the class-A rhodopsin-like family of GPCRs ^30^, whose structure is generally divided into three parts (Fig. 1): an N-terminal extracellular domain (ECD), seven interconnected serpentine transmembrane helices (1-7 TMD) and an intracellular domain (ICD) containing the C terminus ^31^.

Comparison between the sequences of human and carp receptors revealed high similarities (Table 1). Though the percentage of structural similarities between the whole mammalian receptors to the fish receptors was relatively high, ranging from 61% to 67%, the TMD appeared to be much more conserved (75% – 82% similarities), suggesting conserved activation mechanism and function for this domain. Moreover, while cFSHR TMD is more similar to hFSH TMD than to that of hLHR, cLHR TMD is equally similar to both human receptors (Table 1).

**Table 1.** Sequence similarity between carp and human GTHRs. “Whole” was defined as the entire receptor including ECD, TMD extra– and intracellular loops and ICD, whereas “TMD” includes the extra– and intracellular loops that connect the helices.

| Receptors | cLHR |  | cFSHR |  |
| --- | --- | --- | --- | --- |
|  | Whole | TMD | Whole | TMD |
| hLHR | 66% | 75% | 61% | 75% |
| hFSHR | 64% | 75% | 67% | 82% |
| cLHR | 100% | 100% | 60% | 74% |

Traditionally, orthosteric binding sites have been considered the preferred targets for drug development. However, targeting these sites can lead to activation of multiple signaling cascades. For example, the human GTHR can activate internal signal proteins such as Gq/11, Gi/0, IP3, and β-arrestin to regulate various intracellular pathways and mediate receptor internalization, in addition to G proteins and adenylate cyclase pathways. This complexity limits the ability to precisely control synthetic modulation of the receptor ^32^. Despite consistent progress in developing GPCR-targeting allosteric modulators, the enormity of receptor and hormone renders the manipulation of these receptors much more complicated. Hence, only a few small compounds are available for its modulation. We therefore strived to search for candidates that exhibit the potential to directly modulate these receptors. For which, we analyzed *in-silico* the generated structures for possible binding sites and located orthosteric sites on the ECD and allosteric binding sites within the TMD of cGTHRs (Fig. 2; Fig. S1) ^7, 29^.

**Figure 2:**
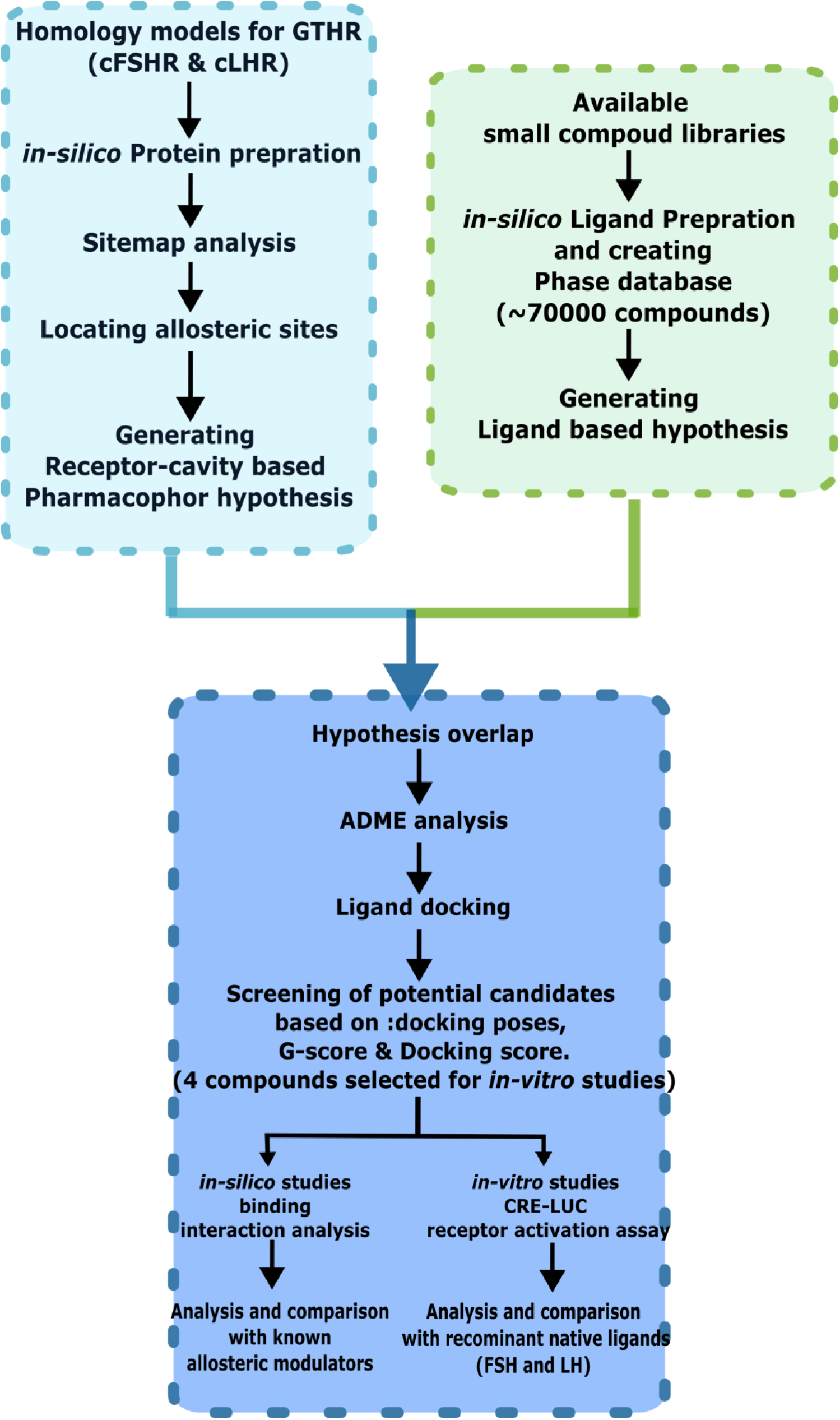
Receptor cavity-based screening procedure for selected compounds.

As the allosteric sites are located close to the ECLs, we hypothesized that the small compounds binding to this site would induce conformational changes similar to those caused by orthosteric binding mechanism. Studies in humans have demonstrated potent allosteric modulation of hFSHR by small compounds, such as Cpd-21f and Org214444-0. These compounds are 10 to 100 times more potent in activating hLHR than in activating hFSHR. The binding site of Cpd-21f and Org-214444-0 almost completely overlapped with that of ligand Org43553 on luteinizing hormone/choriogonadotropin receptor (LHCGR) ^7, 8^. When binding to its allosteric destination on LHCGR receptor, Org43553 was reported to be an almost full agonist, inducing a selective agonistic effect and showing signal cascade specificity ^33^. The allosteric binding pockets in hFSHR and hLHCGR are very similar, and both are mainly composed of residues on TM3, TM5, TM6 and TM7, along with ECL2 and ELC3. The recently published electron microscopy structure of human LHCGR shows Org43553 binding deep in the allosteric pocket at the top half of the TMD (PDB:7FIH), mainly via hydrophobic interactions. Org43553 was reported to be exposed to the hinge domain and ECL ^7^, which induces conformational modulation of the receptor. Based on these findings, we generated a receptor cavity-based pharmacophore hypothesis which is a pharmacophore hypothesis based on the nature of residues on the receptor that are exposed to allosteric binding pockets in the TMD of both cFSHR and cLHR (Fig. 3).

**Figure. 3:**
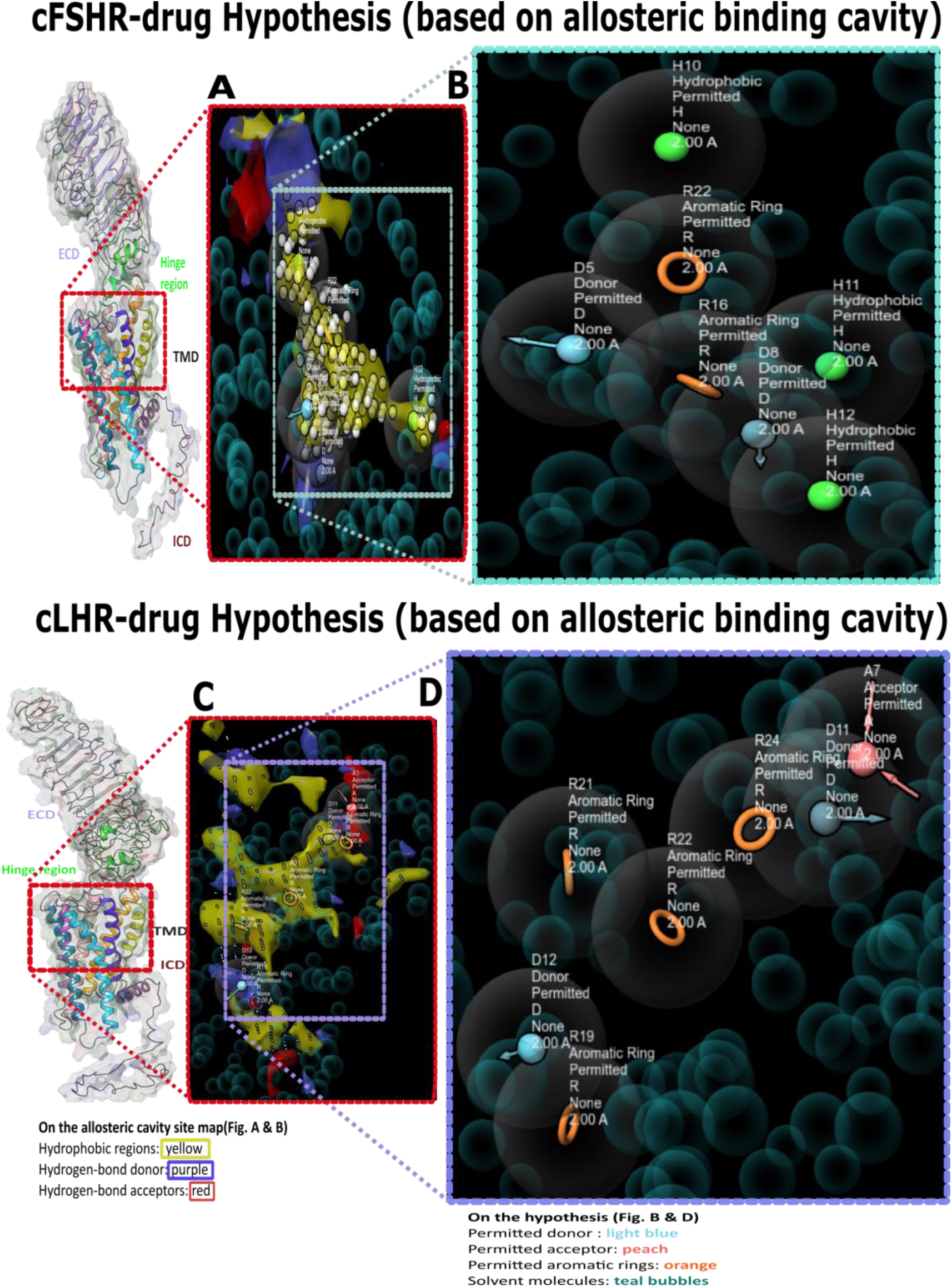
Receptor cavity-based hypothesis map for allosteric modulation. Models showing the surface of exposed residues in the allosteric binding pockets of cFSHR (A) and cLHR (C) and the corresponding receptor cavity-based hypothesis (B, D). Small compounds were screened using these hypothesis maps to select suitable candidate modulators.

GPCR library (54080 compounds https://enamine.net/compound-libraries/targeted-libraries/gpcr-library)) and allosteric GPCR library (14400 compounds https://enamine.net/compound-libraries/targeted-libraries/gpcr-library/allosteric-gpcr-library) were retrieved from Enamine website and were converted to Phase format (see Experimental procedures). We then screened the ligand database based on the generated hypothesis and performed an ADME analysis. We generated ligand-based hypothesis and used them in parallel with the receptor cavity-based hypothesis as an additional screening step and docked the chosen ligands onto the allosteric site. The most successfully docked small compounds were shortlisted based on Glide g-score, XP g-score and Docking score. The docking conformation was selected on the basis of Glide emodel.

Eventually, two compounds were selected for each receptor and tested *in vitro*. These included <u>8028</u> [1-(5-cyclopropyl-2-phenyl-1,3-oxazole-4-carbonyl)piperidin-2-yl]methanamine; <u>4681</u> (1-[2-(2-methylphenyl)-1,3-thiazole-4-carbonyl]piperidin-2-yl)methanamine hydrochloride; <u>9045</u> (1-(2-[(2-chlorophenyl) methoxy] benzoyl) pyrrolidin-3-yl) methanamine; and <u>0801</u> (1-[5-(2,3-dihydro-1,4-benzodioxin-6-yl) –1,3-oxazole-4-carbonyl] pyrrolidin-3-yl) methanamine dihydrochloride) (Figs. 4 and 5).

**Figure. 4:**
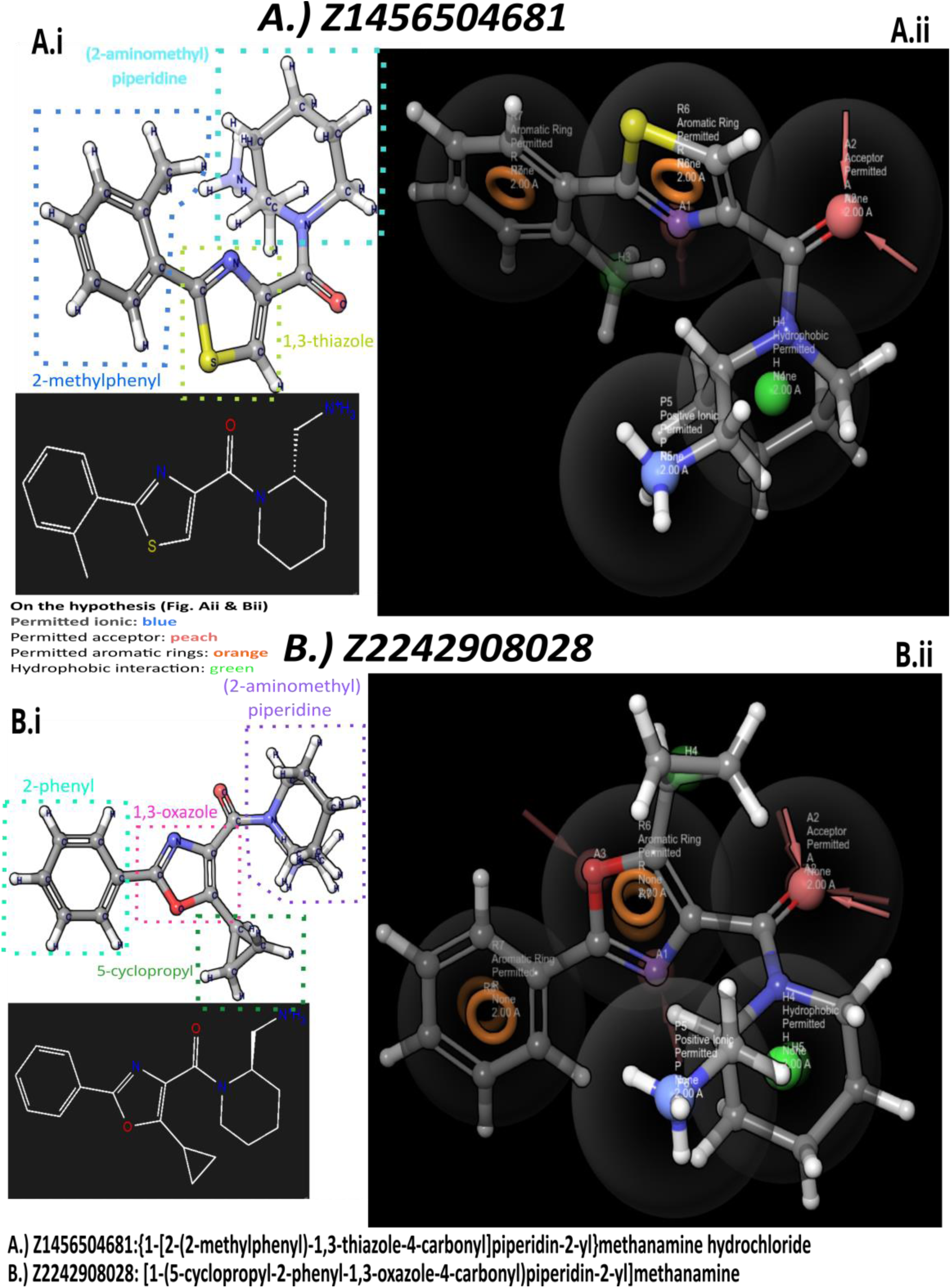
Pharmacophore overlap with the screened ligands Z1456504681 and Z2242908028. **Ai;Bi**: 3D and Kekulé structures of the tested compounds. **Aii,Bii:** Overlap of receptor cavity and ligand based Hypothesis on the small compounds.

**Figure. 5:**
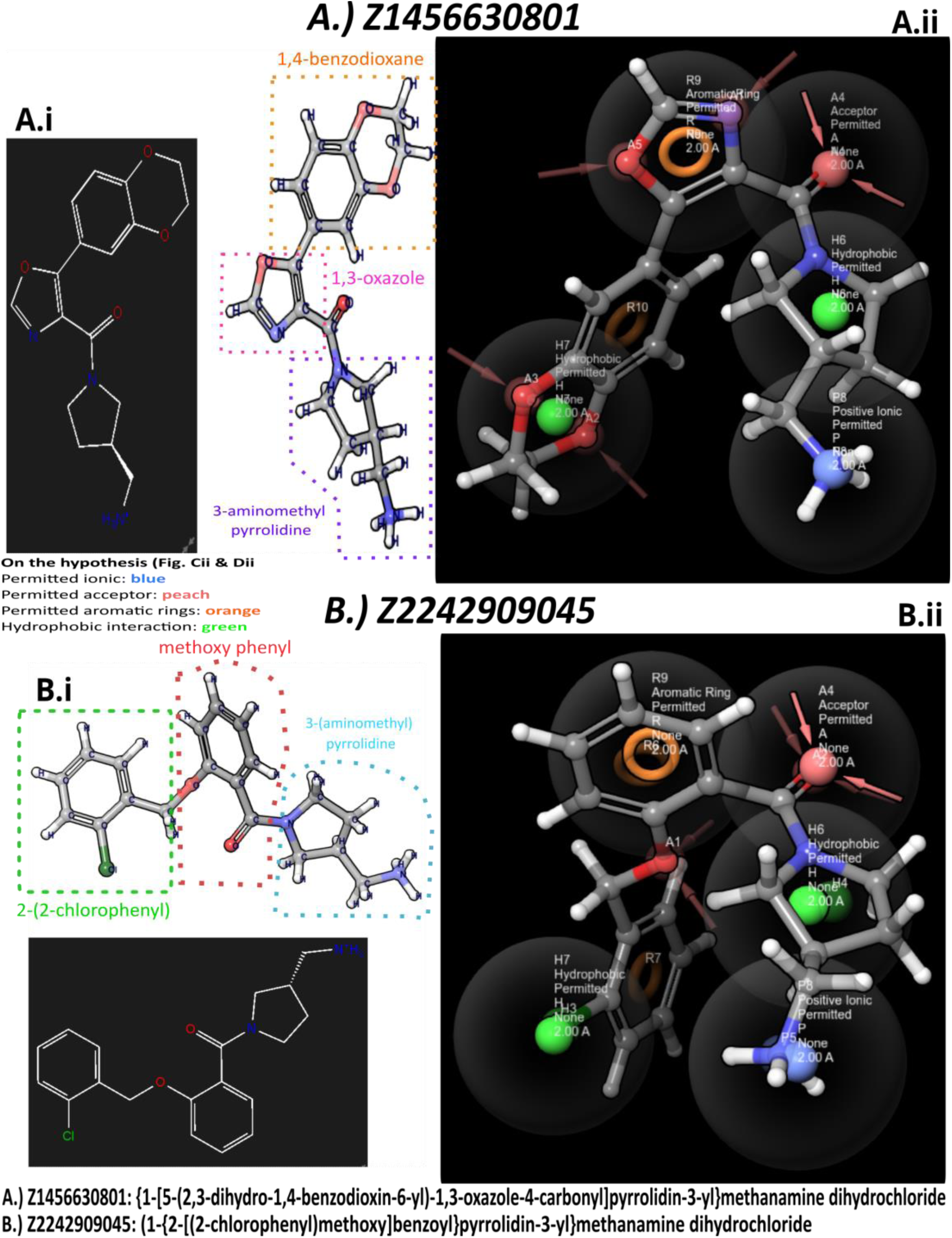
Ligand hypothesis overlap. **Ai;Bi:** 3D and Kekulé structures of the tested compounds. **Aii,Bii:** Overlap of receptor cavity and ligand based HypothesisHypothesis overlap on the small compounds.

The effect of the selected compounds was tested using a receptor transactivation assay on mammalian COS7 cells with cFSHR and cLHR co-transfected together with cAMP response element-luciferase (CRE-LUC), which had been previously shown to be the dominant signal for gonadotropin receptors ^9^. The activity of each compound, as determined by maximum response and EC_50_), was compared to the activity of the recombinant protein previously shown to activate each receptor ^9, 10^. All the four molecules induced agonistic activation of the receptors, albeit with varying response levels and efficiencies toward the different receptor types. The cFSHR was activated by molecules <u>0801</u> and <u>8028</u> (maximum response, 1.406 and 1.499; EC_50_, 23.8 and 4.134 nM, respectively) more efficiently than the recombinant ligands cFSH and cLH at significantly lower EC_50_ values (maximum response, 1.506 and 1.425; EC_50_, 146.5 and 172.8 nM, respectively) (Table 2; Figs. 6 and 7). There was no significant response seen in non GTHR transfected cell lines in response to the selected ligands.

**Fig. 6:**
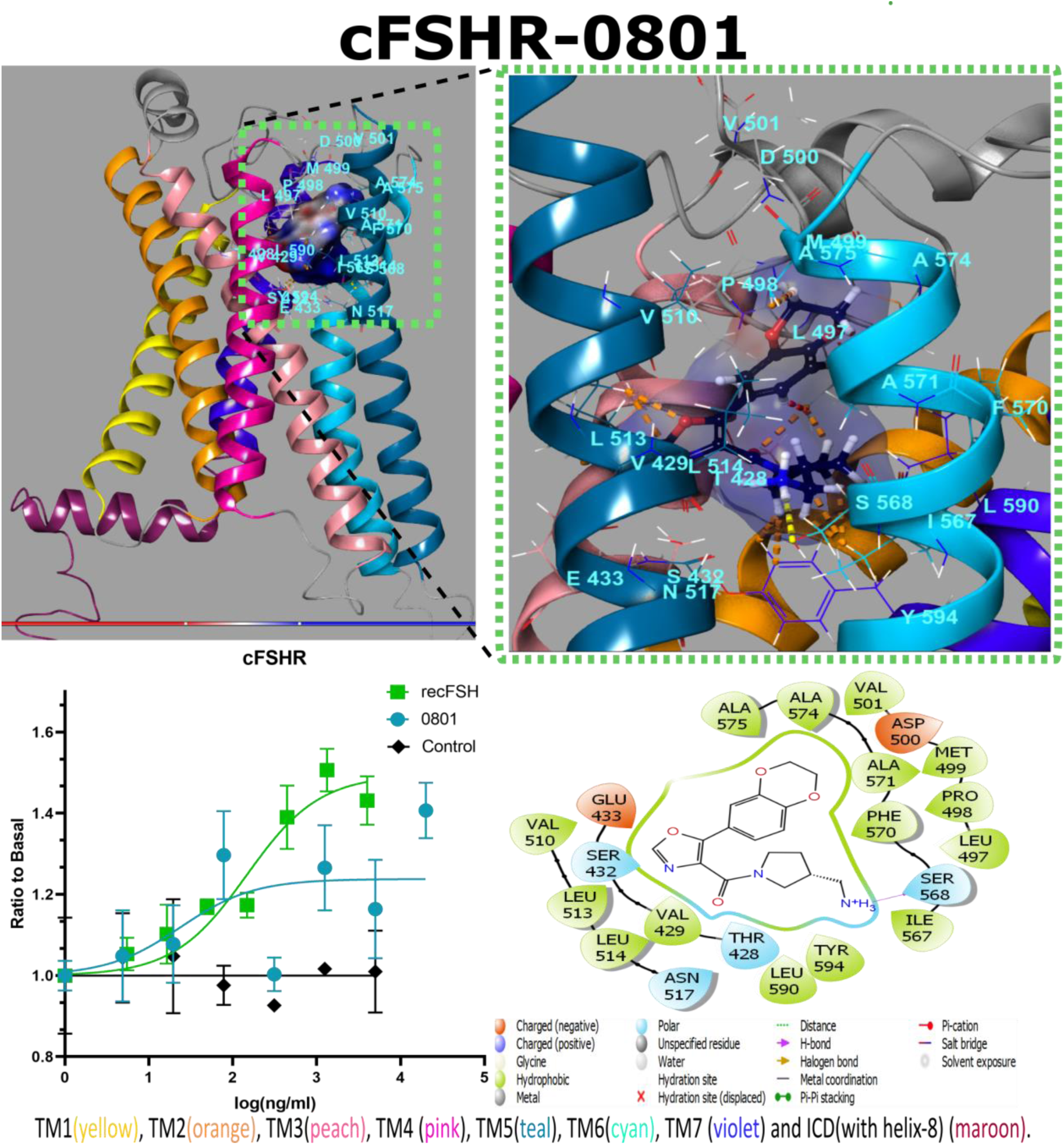
Interactions between cFSHR and Z1456630801. The molecule has 1,3-oxazole in its core, like Z2242908028, instead of dihydroimidazole core on Cpd-21f. This oxazole is attached to 1,4-benzodioxane and 3-aminomethyl pyrrolidine functional groups. On the latter, the aminomethyl sidechain interacts with I567^6.51^ and S568^6.52^ on the TM6 of the receptor via hydrogen bond, whereas L513^5.43^ (TM5) and Y594^7.42^ (TM7) are involved in hydrophobic interactions. Simultaneously, the oxazole interacts with E433^3.37^ (TM3) and L513^5.43^ (TM5). The graph shows CRE-luciferase activity in response to the tested molecule and recombinant carp FSH as a function of concentration. The control denotes to activity of the non-transfected COS 7 cell in response towards concerned ligand.

**Fig. 7.**
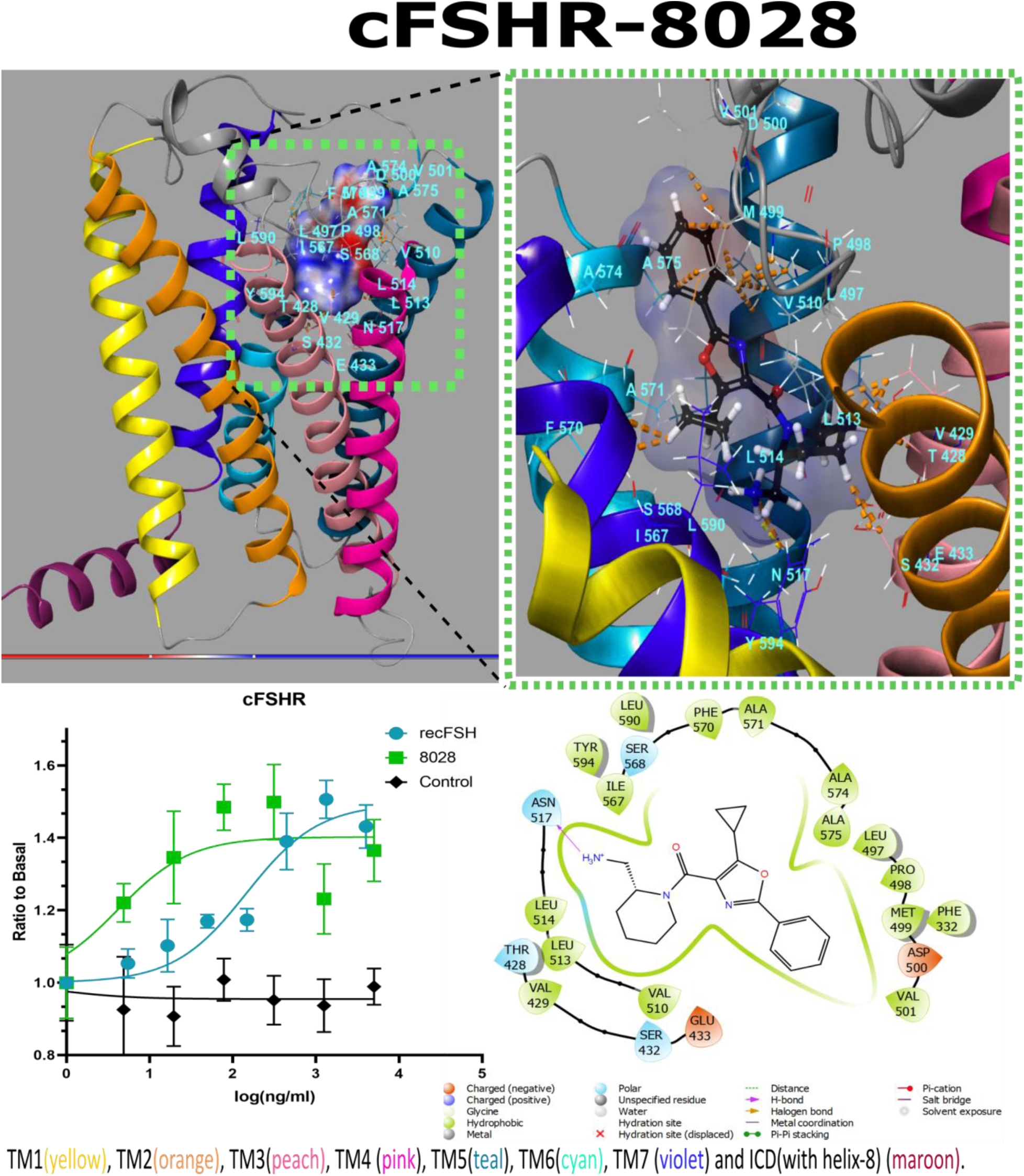
Interactions between cFSHR and Z2242908028. The molecule’s oxazole core is attached to piperidine, phenyl and cyclopropyl sidechains. The phenyl ring was observed to associate with M499 and V501 on ECL2 and with V510^5.40^ on TM5. The cyclopropyl interacts with F570^6.54^ (TM6), whereas 2-aminmethyl piperidine interacts with V429^3.33^ (TM3), S432^3.36^ (TM3), L513^5.43^ (TM5) and N517^5.17^ (TM5). The graph shows CRE-luciferase activity in response to the tested molecule and recombinant carp FSH as a function of concentration. The control denotes to activity of the non-transfected COS 7 cell in response towards concerned ligand.

**Table 2.** EC50 values and activation potency of the tested small compound vs the native ligands. Relative Potency was defined as log (Relative Potency) = log (EC50 of the native compound) – log (EC50 of the novel compound). *Compounds 0801 and 4681 showed potential antagonistic behavior towards cLHR and cFSHR respectively and are marked in red as the ratio does not reflect the overall performance of these small compound.**No significant activity was found by non-transfected Cos7 cells on simulation with the ligands.*

| Ligands | M.W (kDa) | Receptors |  |  |  |  |  |  |  |  |  | Ligand performance |
| --- | --- | --- | --- | --- | --- | --- | --- | --- | --- | --- | --- | --- |
|  |  | cFSHR |  |  |  |  |  | cLHR |  |  |  |  |
|  |  | *EC<sub>50</sub>* (nM) | *EC<sub>50</sub>* (ng/ml) | *log EC<sub>50</sub>* (ng/ml) | *Max. Response* | *log (Relative Potency)* | *EC<sub>50</sub>* (nM) | *EC<sub>50</sub>* (ng/ml) | *log EC<sub>50</sub>* (ng/ml) | *Max. Response* | *log (Relative Potency)* |  |
| *recFSH* | 25 | 5.86 | 146.5 | 2.166 | 1.50655 | 0 | 0 | - | - | - | - | cFSHR specific native hormone |
| *recLH* | 30 | 5.76 | 172.8 | 2.482 | 1.42585 | -0.316 | 10.11 | 303.3 | 0 | 1.21619 | 0 | cFSHR & cLHR native hormone |
| 9045 | 0.344835 | 9882.988 | 3408 | 3.532 | 1.30924 | -1.366 | 26937.52 | 9289 | -1.485 | 1.39297 | -1.485 | High agonist for cFSHR & cLHR |
| 0801* | 0.32935 | 72.26355 | 23.80 | 1.377 | 1.40680 | 0.789 | 16.49917 | 5.434 | 1.7468 | 1.03843 | 1.7468 | cFSHR specific agonist and potential antagonist for cLHR |
| 8028 | 0.325405 | 12.70417 | 4.134 | 0.6164 | 1.49917 | 1.5496 | 1876.124 | 610.5 | -0.304 | 1.21069 | -0.304 | cFSHR & cLHR selective agonist (partial towards cFSHR) |
| 4681* | 0.315433 | 97.77037 | 30.84 | 1.489 | 1.22959 | 0.677 | 300.6 |  | 0.004 | 1.16487 | 0.966406 | cLHR specific agonist and potential antagonist for cFSHR |

cLHR was activated by molecules 8028 and 4681 (max response, 1.210 and 1.164; EC_50_, 610.5 and 300.6 nM, respectively) at similar levels as the recombinant cLH (max response, 1.216; EC_50_, 303.3 nM) (Table 2). Therefore, we report molecules <u>4681</u> and <u>0801</u> as specific agonists of cLHR and cFSHR, respectively. Moreover the 4681 appear to be potential antagonist for FSHR and <u>0801</u> for cLHR. Molecule 8028 is a dual agonist for both receptors; however, it activated cFSHR at a significantly lower dose. Regarding molecule <u>9045</u>, although it also activated both receptors, the EC_50_ values reflect a very low efficiency compared to the recombinant ligands. Despite both 4681 and 0801 showing potential antagonistic behaviour towards the cFSHR maximal response at some doses spiked, this might be the result of constitutive activity of the receptor that remain unblocked by the antagonist. (Table 2).

The <u>0801</u>, which specifically activated cFSHR, bound to an allosteric binding site that is positioned similarly to the binding sites reported for Cpd_21f-cLHR and Org43553-cFSHR interactions in human homolog ^7, 8^; however, it interacted with the lower region of the binding pocket, majorly via hydrophobic interactions (Figs. 4ii and 6). The 1,4-benzodioxane group is exposed to the cFSHR ECL2 and interacts with M499_cFSHR_, but simultaneously it also showed interactions with A571^6.55^ and A575^6.59^. *In silico* analysis showed that various substitution mutations in I567^6.51^_cFSHR_ and A571^6.55^ on TM6 caused the most significant decrease in complex stability and ligand affinity; thus, these residues may play a key role in receptor activation. Mutations in similarly positioned homologs in hLHCGR (I585^6.51^W_hLCGHR_ and A589^6.55^F_hLCGHR_) have been reported to reduce the ability of Org43553 to activate the receptor. cLHR was not activated in response to <u>0801</u> (Fig. S2) and its activity even slightly decreased with increasing dosses, suggesting this molecule as a potential NAM/NAL for cLHR.

The compound <u>8028,</u> which was more partial towards cFSHR, has an oxazole at its core attached to piperidine, phenyl and cyclopropyl sidechains (Fig. 4Bi, Bii). The phenyl ring interacts with ECL2, which functions as an intramolecular modulator of the TMD.

In vitro results along with structural interactions suggests that the interaction between M499_cFSHR_ and V501_cFSHR_ on the ECL2 of cFSHR is crucial for receptor activation, as these interactions were observed in the <u>8028</u> binding but not in <u>4681</u> (Fig. S3), which did not activate the receptor. The activation of cLHR in response to <u>8028</u> was similar to the response to its native ligand cLH. Our studies show that docking of <u>8028</u> to cLHR occurred comparatively deeper within the allosteric binding cavity of the receptor (Figs. 5i and 7). The phenyl ring attached to the oxazole is also exposed to ECL2 and interacts particularly with M534_cLHR_, P533_cLHR_-, which are positioned similarly to M499_cFSHR_ and V501_cFSHR_ on cFSHR. *In silico* mutation analysis showed that substituting homologous cLHR residues L532_cLHR_, P533_cLHR_ and M534_cLHR_ with various amino acids considerably reduced binding stability and affinity. Despite the conformational variance, the interacting amino acids on cFSHR and cLHR are significantly conserved.

The ECL2 is the largest intracellular loop of both cFSHR and cLHR. Studies in hFSHR have established that ECL2 is indispensable in mediating post-docking conformational changes by interacting with other ECLs and TMDs. The homologous mutation P519T_hFSHR_(P533_cLHR_), which is positioned on hFSHR-ECL2, has been associated with primary amenorrhea in patients, whereas V514A_hFSHR_ (V501_cFSHR_) mutation was observed in patients undergoing *in vitro* fertilization who exhibited symptoms of iatrogenic ovarian hyperstimulation syndrome. Further, the P519T mutation on hFSHR ECL2 ultimately impaired adenylate cyclase stimulation *in vitro* ^34, 35^. P519_hFSHR_ is highly conserved in hFSHR(P516_hLHCGR_), cFSHR(P498_cFSHR_) and cLHR(P533_cLHR_), and its mutation was reported to disrupt receptor trafficking to the cell surface and subsequently abolished FSH binding and cAMP production ^29^. Therefore, the interaction of the ligand with this residue might explain its agonistic effect on cLHR and cFSHR. Further, whereas F515^ECL2^A_hLHCGR_ (homologs L532^ECL2^_cLHR_, L497^ECL2^ and L518^ECL2^_hFSHR_) and T521^ECL2^A_hLHCGR_ (homologs L538 ^ECL2^, L503 ^ECL2^ and L524 ^ECL2^) mutations on hLHR ECL2 enhanced internalization and cAMP signaling, S512A_hLHR_ (S529_cLHR_, F515_hFSHR_), and V519A_hLHR_(homologs I536^ECL2^_cLHR_, V501^ECL2^_cFSHR_ and I522^ECL2^_hFSHR_) impaired these processes ^36^. This indicates that ECL2 might play a key role in selective activation of downstream signal transduction and impact its efficiency significantly and is therefore a potential target for signaling pathway-specific selective modulators.

The cLHR ELC2 showed interactions with the hinge domain and ECL1, and *in silico*, mutations in L532A_cLHR_, P533A_cLHR_, M534A_cLHR_ on ECL2 were seen to substantially reduce complex stability and ligand affinity (Fig. 8). The oxazole was also observed to interact with A606^6.55^_cLHR_ on TM6, which seems to play a crucial role in <u>8028</u> binding. *In silico* mutations analysis showed that substitution of cLHR A606^6.55^ with various amino acids (A606F_cLHR_, A606D_cLHR_, A606R_cLHR_, A606W_cLHR_) caused the most significant decrease in complex stability and ligand affinity. Studies have shown that mutation in the similarly positioned hLHCGR residue A589^6.55^W_hLHCGR_ has reduced activation by Org43553 ^7^, suggesting its role in allosteric receptor activation. As <u>8028</u> docking occurs comparatively deeper inside the allosteric TMD binding pocket of cLHR, it shows many more interactions with the TM helices than when docking onto cFSHR. We hypothesize that these interactions might hamper the post-binding conformational changes, therein reducing the activation of cLHR by <u>8028</u> compared to cFSHR.

**Fig. 8:**
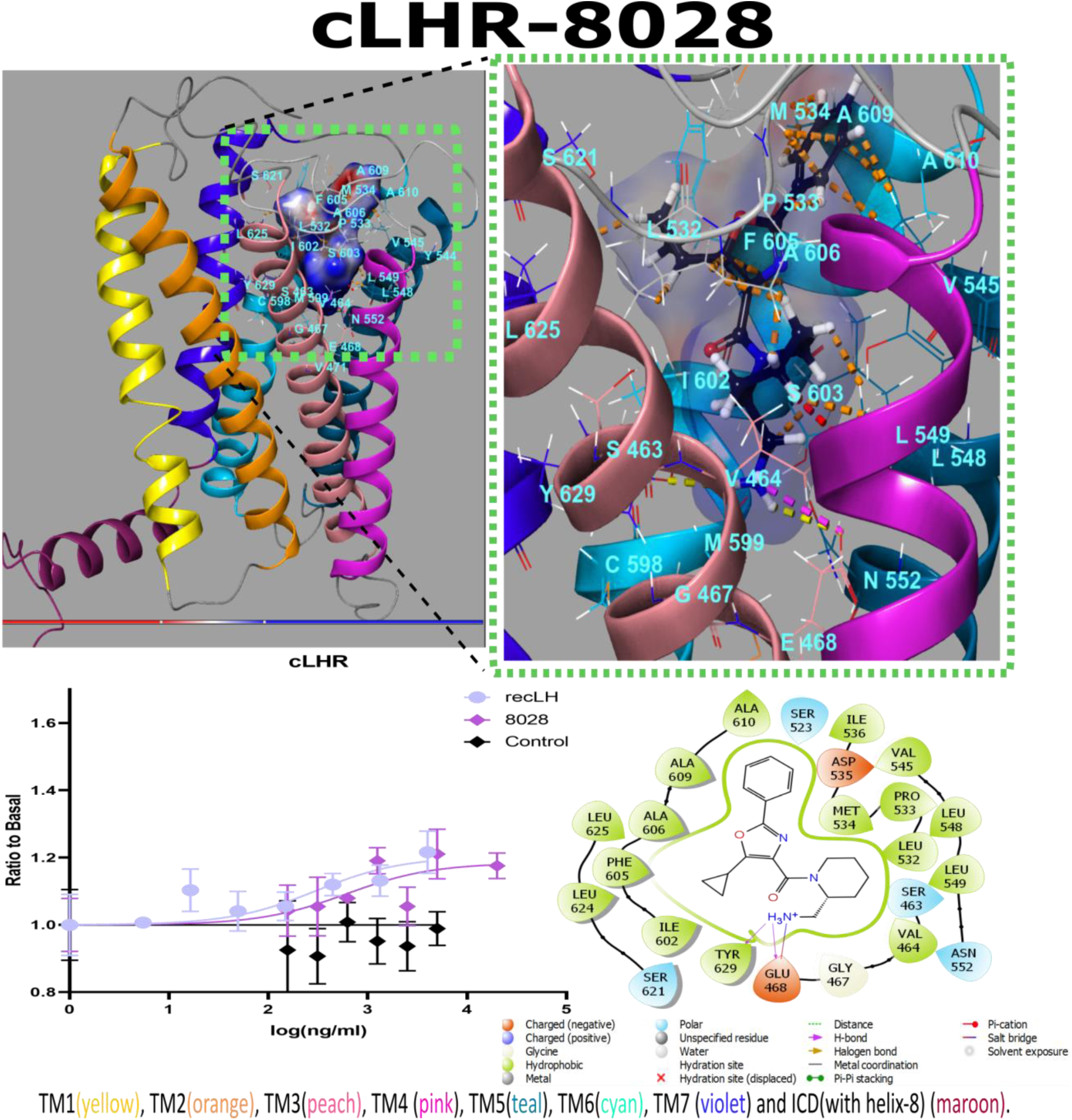
Interactions between cFSHR and Z2242908028. This compound docked comparitively deeper into the allosteric cavity. Both the 1,3-oxazole core and the attached 5-cyclopropyl and (2-aminomethyl) piperidine sidechain closely interacted with L532 on ECL2, whereas the phenyl ring interacted with M534 and P533 on ECL2. Major interactions were observed with hydrophobic residues on TM5 (V545^5.40^, L548^5.43^ and L549^5.44^) TM6 (A606^6.55^ and A609^6.58^) and TM7 (L625^7.38^). The graph shows CRE-luciferase activity in response to the tested molecule and recombinant carp LH as a function of concentration. The control denotes to activity of the non-transfected COS 7 cell in response towards concerned ligand.

The compound <u>4681</u> induced a similar response in cLHR activity as did <u>8028</u>, which also has a similar structure. However, the oxazole core of <u>8028</u> is replaced by 1,3-thiazole in <u>4681,</u> and the cyclopropyl side chain is absent (Fig. 4i). Another difference is the presence of a 2-methylphenyl group attached to the thiazole core, instead of a phenyl ring. This bulkier functional group leans more backward towards the TM7, while the methyl extension interacts with F605^6.54^ on cLHR TM6. At the opposite end, the phenyl ring simultaneously associates with L532 on ELC2, which might slightly restrict TM6 movement (Fig. 9), whereas the shorter phenyl group of <u>8028</u> binds to A609^6.58^_cLHR_ and M534^ELC2^ instead. Although the cyclopropyl sidechain lacks interactions with the 2-aminomethyl piperidine group, Y629 seems to compensate for the lack of interaction with L625^7.38^ positioned on TM7, which appears to play a crucial role in the conformational modulation of TM7. Moreover, (2-aminomethyl) piperidine interacts with Y629, which is situated comparatively much deeper and hence might impair TM7 movement, further reducing the activation potential of <u>4681</u>. This might explain the lower potency values observed in response to this molecule (Table 2). Although <u>4681</u> bound to cFSHR, the observed receptor activity was significantly lower than cFSH. The residue mostly interacted with the deep hydrophobic pocket created by the TM helices and showed no contact with the ECL2, which we hypothesize is crucial for allosteric site-mediated docking and signal pathway specificity. The cFSHR, being promiscuous, is activated not only by its cognate ligand cFSHR, but also by cLHR^9^. The compound showed strong interactions with M499_cFSHR_, L513^5.43^, I567^6.51^ and A571^6.55^_cFSHR_, which seem crucial for cFSHR activation. With its considerably low receptor activation, <u>4681</u> has substantial potential as a cFSHR NAL/NAM or a cLHR-specific allosteric agonist.

**Fig. 9:**
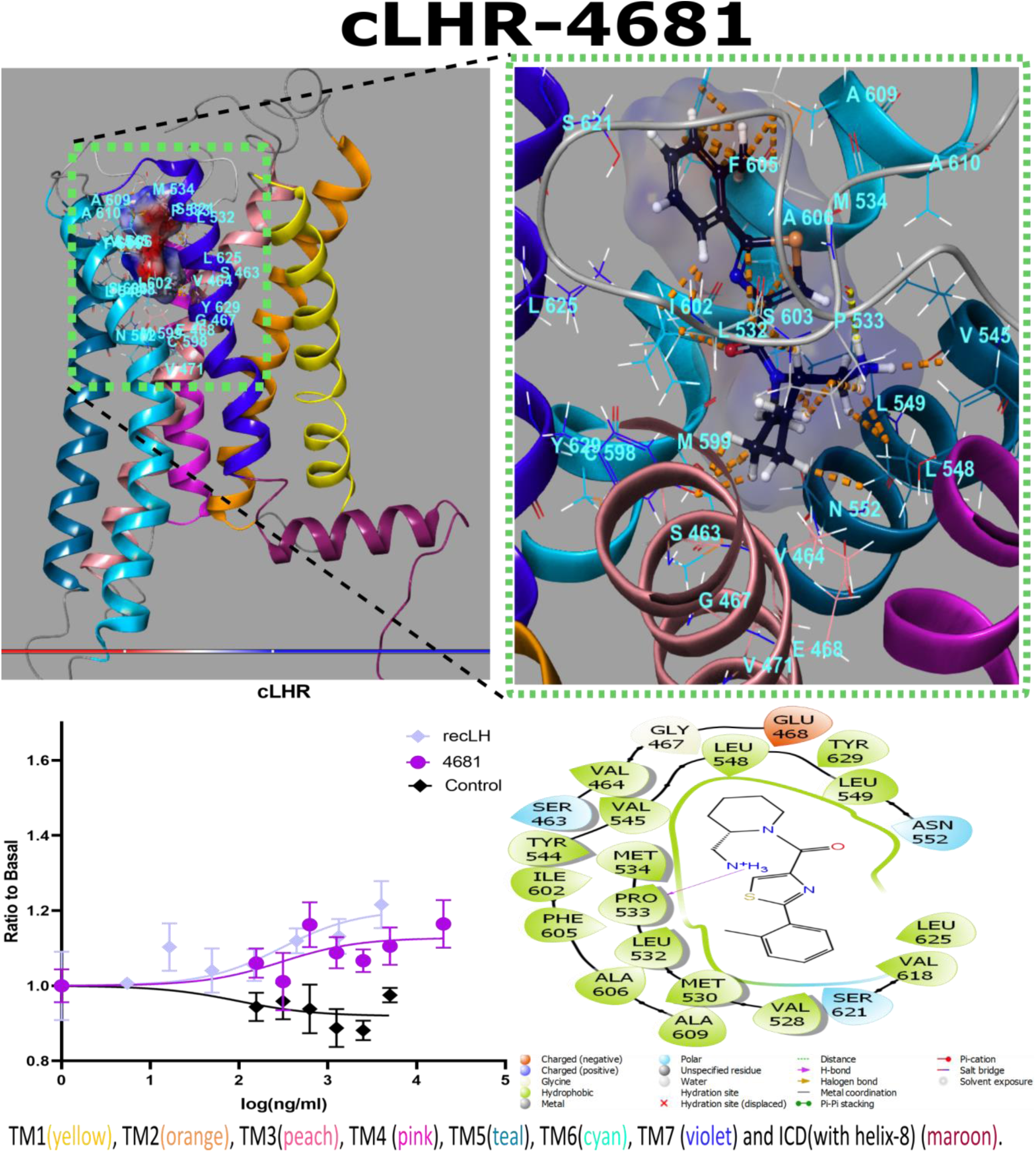
Interactions between cLHR and Z1456504681. The molecule has 1,3-thiazole at its core, which is attached to piperidine, 2-methylphenyl and cyclopropyl sidechains. The (2-aminomethyl) piperidine sidechain closely interacted with L532 and P533 on ECL2. Other key residues exposed to the functional groups include M534 on ELC2, V545^5.40^, L548^5.43^ and L549^5.44^ on TM5; F605^6.54^ and I602^6.51^ on TM6; and Y629^7.42^ on TM7. The graph shows CRE-luciferase activity in response to the tested molecule and recombinant carp LH as a function of concentration. The control denotes to activity of the non-transfected COS 7 cell in response towards concerned ligand.

The 9045 induced receptor activation at much higher concentrations than both 8028 and recFSH, but a gradual dose-dependent increase was observed. Its docking conformation notably differed between binding to cFSHR and cLHR. The molecule has a methoxyphenyl core that is attached to 2-(2-chlorophenyl) and 3-(aminomethyl) pyrrolidine functional groups (Figs. 5ii and 7). Org214444-0 has a similar oxyphenyl core, which is approximately twice as bulky due to the functional groups attached to it. 9045 docked to cFSHR in a horseshoe conformation. Although it interacted with L497_cFSHR_, P498_cFSHR_ and M499_cFSHR_ on ECL2, there were many interactions observed with the TM helices, e.g., with I567^6.51^_cFSHR_ (TM6), S568^6.52^_cFSHR_ (TM6), A571^5.55^_cFSHR_ (TM6), N517^5.47^_cFSHR_, and Y594^7.42^_cFSHR_ (TM7).

Various in silico mutations in the key residues A571^6.55^, and I567^6.51^, as well as in P498_cFSHR_ and M499_cFSHR_ on the ECL2, significantly decreased the affinity and stability of the cFSHR-9045 complex. When binding to cLHR, 9045 displayed a much more linear conformation than upon binding to cFSHR, forming mainly hydrophobic interactions and binding deeper into the binding pocket (Fig. S4). The ligand penetrated more deeply into the hydrophobic cavity as compared to 8028. Similar to its docking to cFSHR, the 3-(aminomethyl) pyrrolidine functional group engaged with surrounding TM helices at E468^3.37^ (TM3), N552^5.47^ (TM5), I602^6.51^_cLHR_ (TM6) and Y629^7.42^_cLHR_ (TM7), which played a crucial role in post-docking conformational modulation of cLHR (Fig. S5). Further, upon binding to 9045, the orientation of cLHR residues I602^6.51^ and N552^5.47^ and cFSHR residues Y594^7.42^ and N517^5.47^ in cFSHR diverged towards the ligand and internally engaged other exposed residues on the surrounding TM helices. We hypothesize that these interactions of 9045 with both cFSHR and cLHR substantially restrict the post-docking conformational changes, hence accounts for the lower potency of the molecule.

The in-silico method we used to generate receptor cavity-based hypothesis and for pharmacophore screening considerably increases the probability of identifying small compounds capable of receptor binding and pathway-specific modulation. Though this approach has been developed for efficient pharmacophore design, a few studies have tested with potential GPCR modulators, and none have focused on GTHRs. Due to the elaborate activation mechanism of its extracellular domain, GTHR activation through orthosteric binding of its cognate receptor is elaborate and complicated. Moreover, piscine GTHR lacks the strict hormone-receptor specificity seen in mammals, as both FSHR and LHR are variably promiscuous, depending on the fish species.

Activation can induce variable signal transduction cascades in response to the same stimulation due to the large size of both receptor and hormone, which may form complexes larger than 1000 amino acid-long. Irregularities or mutations in these molecules may lead to various reproductive disorders. Allosteric binding sites provide an alternative route for activation and modulation of GTHRs, which may offer more efficient regulation of downstream signaling cascades. Screening and selection of small compound modulators are expensive and time-consuming, with low return of successful hits. Our in vitro analyses showed the efficiency of using a receptor cavity-based hypothesis for in silico screening of small compounds with agonist effects. The molecules can be further improved or turned into potential NAMs/NALs by replacing the functional groups attached to the pharmacophore core. Overall, allosteric sites in GTHRs show great potential for receptor manipulation while bypassing the elaborate ECD-based orthosteric activation mechanism. While allosteric modulators can act in a regulatory capacity as PAMs, NAMs and NALs, they can also directly manipulate the TMD independently of orthosteric mechanisms. Thus, they may be a crucial tool to overcome the lack of post-binding signaling specificity seen in orthosteric GTHR activation.

## Conclusion

Maintaining a controlled reproductive cycle of fish is of utmost importance in aquaculture. Currently, most species commonly use hormonal manipulation to regulate gonadal activity. However, these hormonal treatments often have limitations, such as high costs and limited effectiveness. To overcome these challenges, there is significant potential in utilizing allosteric modulators as regulators of hormonal activation.

Our research employed the receptor cavity-based hypothesis and ligand screening method to identify allosteric agonists capable of activating receptors independently from native ligands. Through this approach, we successfully selected four small compounds as potential modulator drug candidates for cyclic gonadotropin-releasing hormone receptors (cGTHRs). Our novel pharmacophore screening procedure, which incorporated multiple in silico screening stages, ADME (absorption, distribution, metabolism, and excretion) considerations, and docking results, significantly enhanced the efficiency of the screening process. The efficacy of our selected compounds was further confirmed through in vitro testing.

Considering the complexity of piscine GTHR-GTH interactions, and the significance of controlling and manipulating fish reproductive cycles, our strategy holds promise for identifying additional allosteric modulators. This approach has the potential to revolutionize the field of aquaculture by providing cost-effective and efficient methods for regulating fish reproduction.

## Data and Software Availability

The homology modelling structure of the inactive GnRH1R structure Were generated using I-TASSER (https://zhanggroup.org/I-TASSER/). The template for homology modelling were obtained from the Protein Data Bank (RCSB PDB: https://www.rcsb.org/). In this work, the site map analysis, hypothesis generation, molecular docking was performed using (Maestro, Schrödinger, LLC, New York, NY, 2021.) and can be downloaded from https://www.schrodinger.com/. The compound libraries can be downloaded from https://enamine.net/compound-libraries. The downloaded libraries were further processed and used to construct conformational databases for ligands using the Phase module (Phase, Schrödinger, LLC, New York, NY, 2021.) and are made available at https://zenodo.org/record/8120822 along with input files receptor homology models, docking grids, and docked ligand receptor complexes. The selected compounds can be ordered from Enamine online store (https://new.enaminestore.com/).

## AUTHOR INFORMATION

### Authors’ contributions

I.A. did the *in-silico* modelling, screening, and analysis. *In vitro* LUC assays and statistical analysis were performed by L.H. H.L. ran control experiments on non-transfected cell lines. The project supervision and arrangement of funding was done by B.L.S. The manuscript was written through contributions of all authors. All authors have given approval to the final version of the manuscript.

### List of Abbreviations

GTH, gonadotropin; GTHR, gonadotropin receptors; cFSH, carp follicle stimulating hormone; cFSHR, carp follicle stimulating hormone receptor; cLH, carp luteinizing hormone; cLHR, carp luteinizing hormone receptor; LUC, luciferase activation assay; EC_50_, half maximal effective concentration.

## Funding

This project has received funding from the European Union’s Horizon 2020 research and innovation programme under the Marie Sklodowska-Curie Grant Agreement no. 642893 – IMPRESS and the USDA, NIFA-BARD Research Project IS-8339-23.

**Figure. S1:**
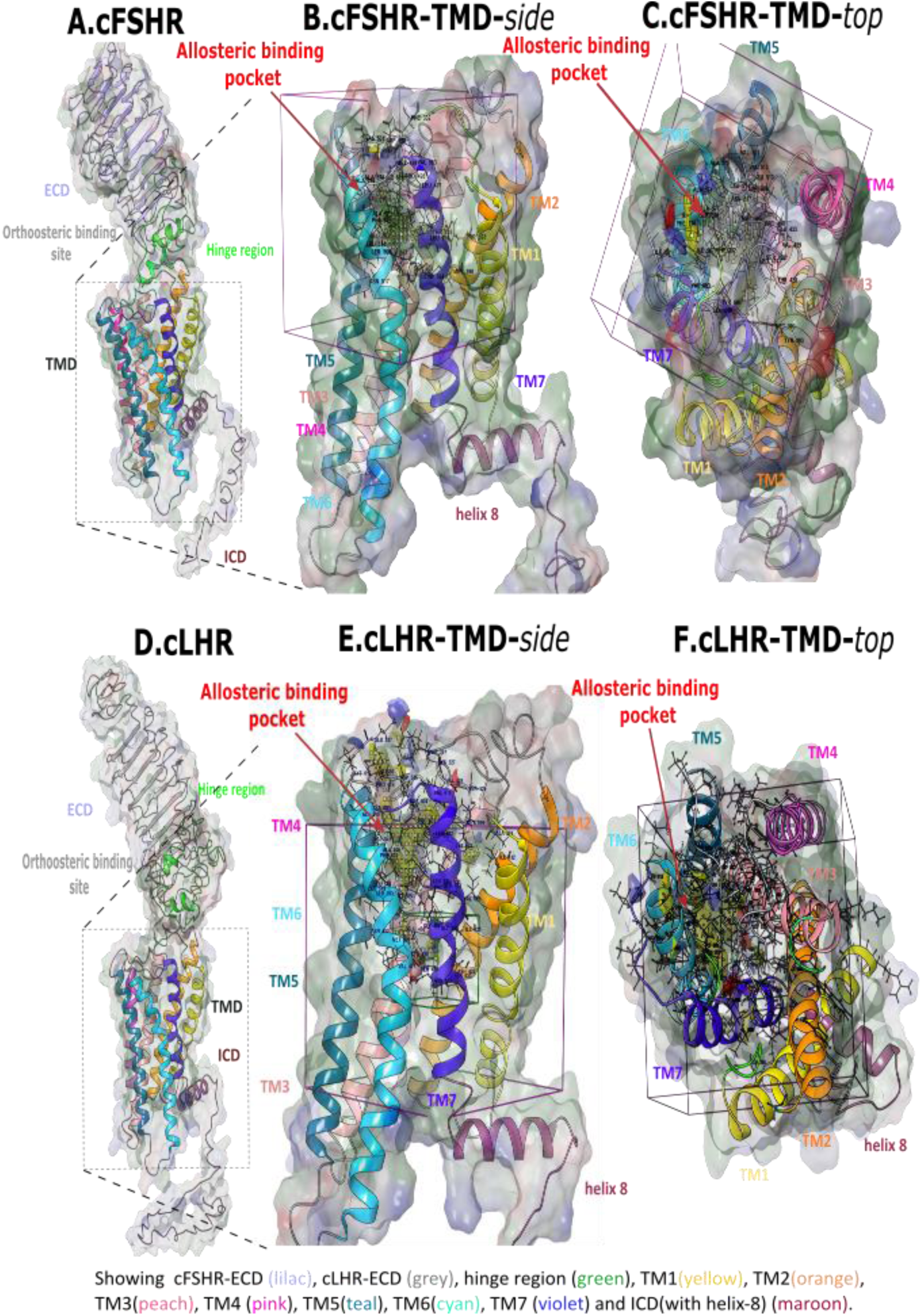
Ribbon diagrams showing the targeted allosteric binding sites located within hydrophobic transmembrane cavities in cFSHR (A-C) and cLHR (D-E). The transmembrane regions of cFSHR are magnified in B (side view) and C (top view), and of cLHR in E (side view) and F (top view).

**Figure. S2.**
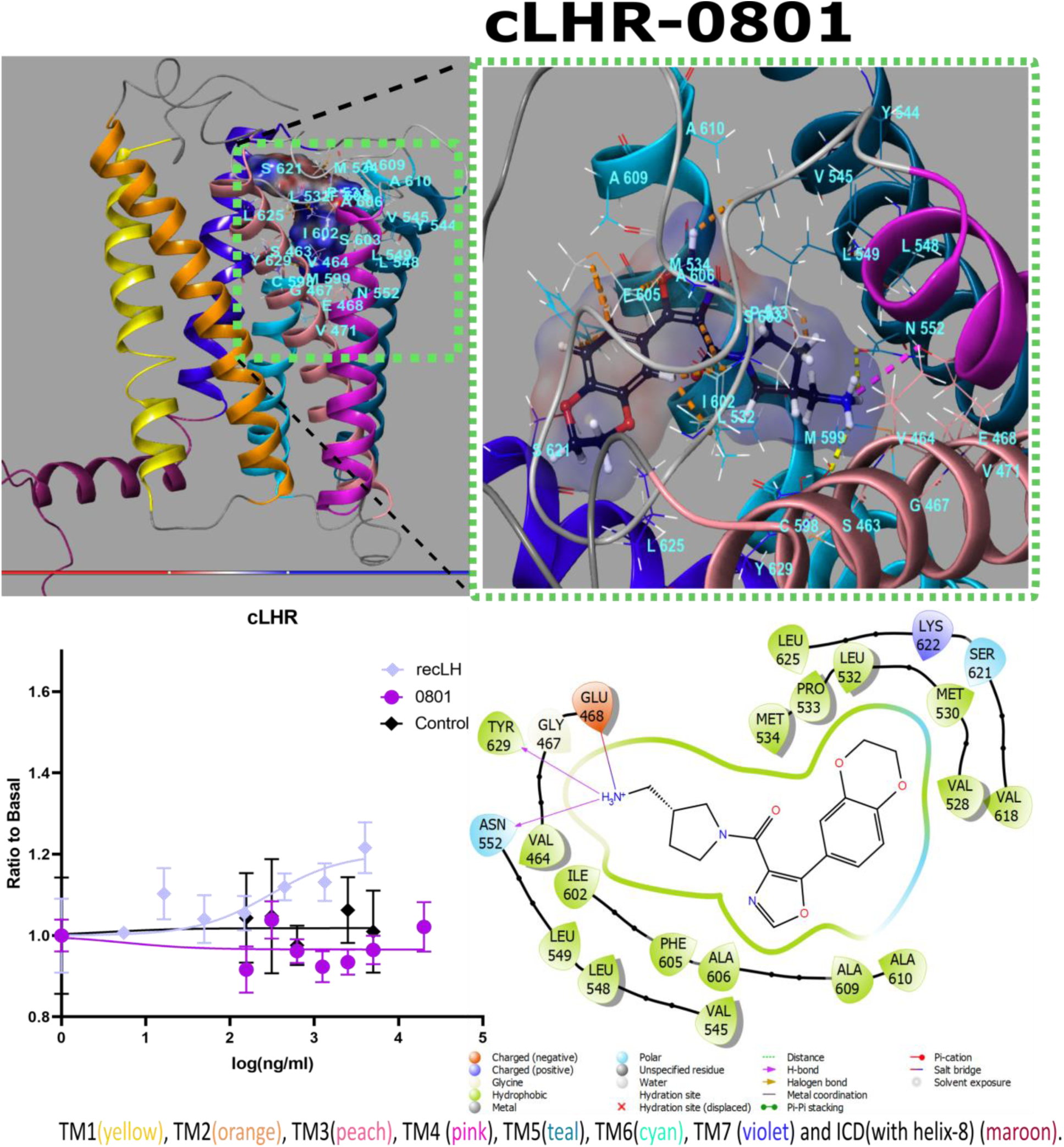
Interactions between cLHR and Z1456630801. The graph shows CRE-luciferase activity in response to the tested molecule and recombinant ligand as a function of concentration. The control denotes to activity of the non-transfected COS 7 cell in response towards concerned ligand.

**Fig. S3.**
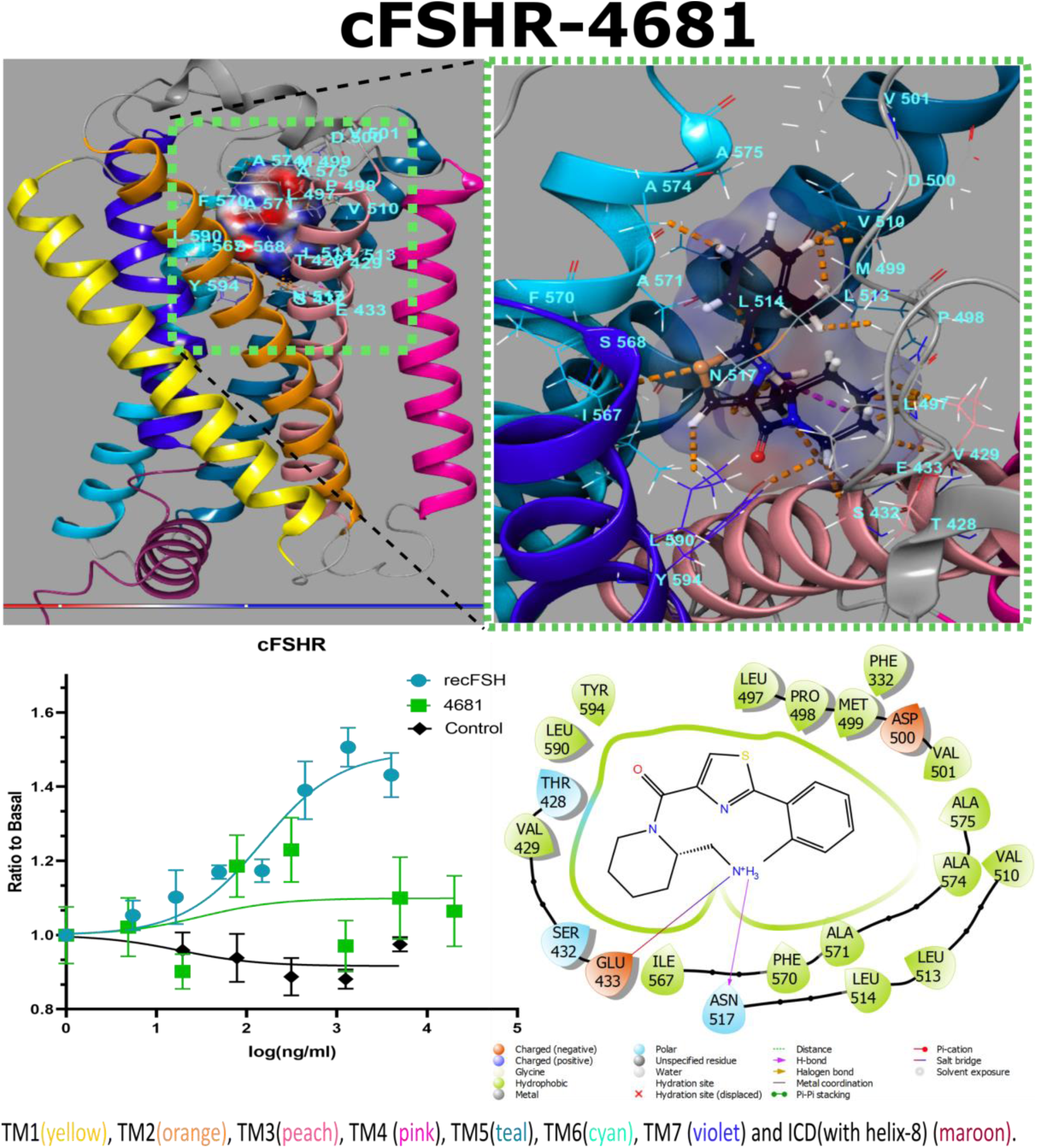
Interactions between cFSHR and Z1456504681. The graph shows CRE-luciferase activity in response to the tested molecule and recombinant ligand as a function of concentration. The control denotes to activity of the non-transfected COS 7 cell in response towards concerned ligand.

**Figure. S4.**
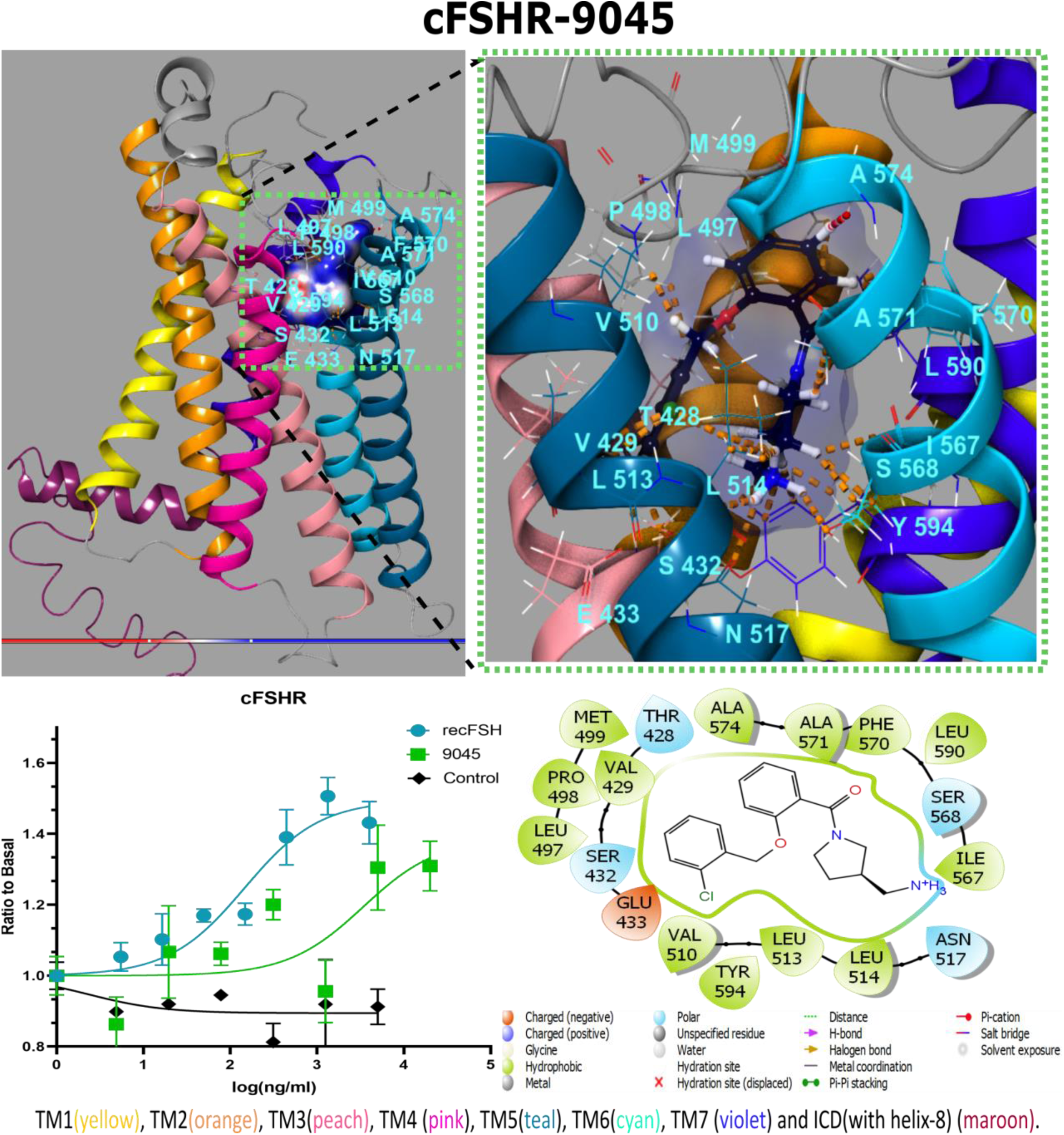
Interactions between cFSHR and Z2242909045. The molecule has a methoxyphenyl core that is attached to 2-(2-chlorophenyl) and 3-(aminomethyl) pyrrolidine functional groups. The chlorophenyl group interacted with L497 and P498 on ECL2, L513^5.43^ and L514^5.44^ on TM5, and with V429^3.33^ and S432^3.36^ on TM3. The methoxyphenyl group showed associations to M499 (ECL2) and the aminomethyl pyrrolidine group interacted with I567^6.51^, S568^6.52^ and A571^5.55^ (TM6), Y594^7.42^ (TM7, internally interacts with S432^3.36^ on TM3), L514^5.44^ (TM5), and N517^5.47^ (TM5; interacts with E433^3.37^ on TM3 internally). The graph shows CRE-luciferase activity in response to the tested molecule and recombinant ligand as a function of concentration. The control denotes to activity of the non-transfected COS 7 cell in response towards concerned ligand.

**Fig. S5.**
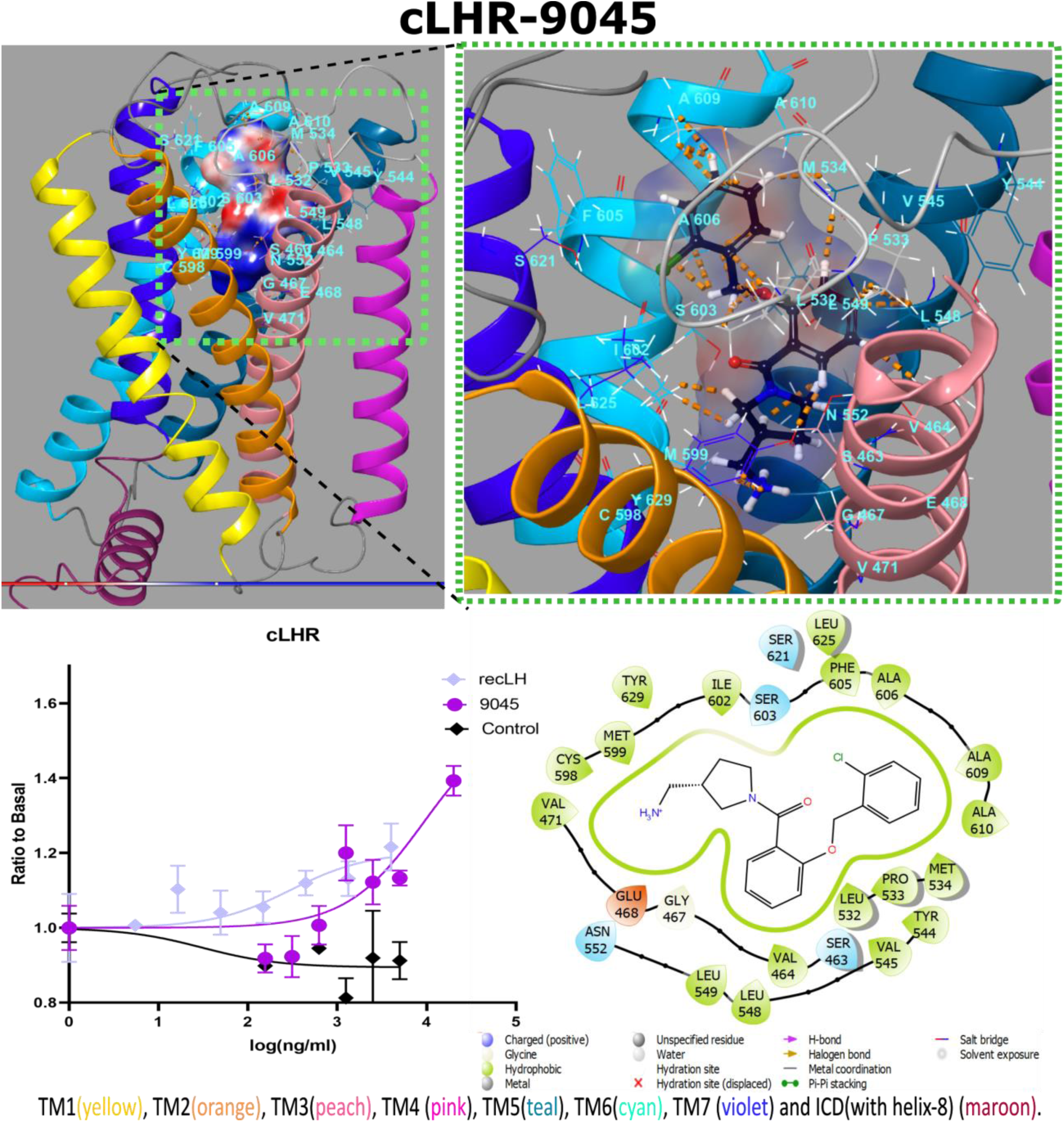
Interactions between cLHR and Z2242909045. The chlorophenyl group is exposed to ECL2 and interacts mainly with L532 and M534 and also with A606^6.55^ and A609^6.58^ (TM6). The oxyphenyl core interacts with P533 on the ECL2 and with V464^3.33^ (TM3), V545^6.55^, L548^5.43^ and L549^5.44^ (TM5), and Y629^7.42^ (TM7). The aminomethyl pyrrolidine group interacts with I602^6.51^ (internally interacts with M599^6.48^), N552^5.47^ (internally interacts with E468^3.37^), and Y629^7.42^ (TM7). The graph shows CRE-luciferase activity in response to the tested molecule and recombinant ligand as a function of concentration. The control denotes to activity of the non-transfected COS 7 cell in response towards concerned ligand.

## For Table of Contents Use Only

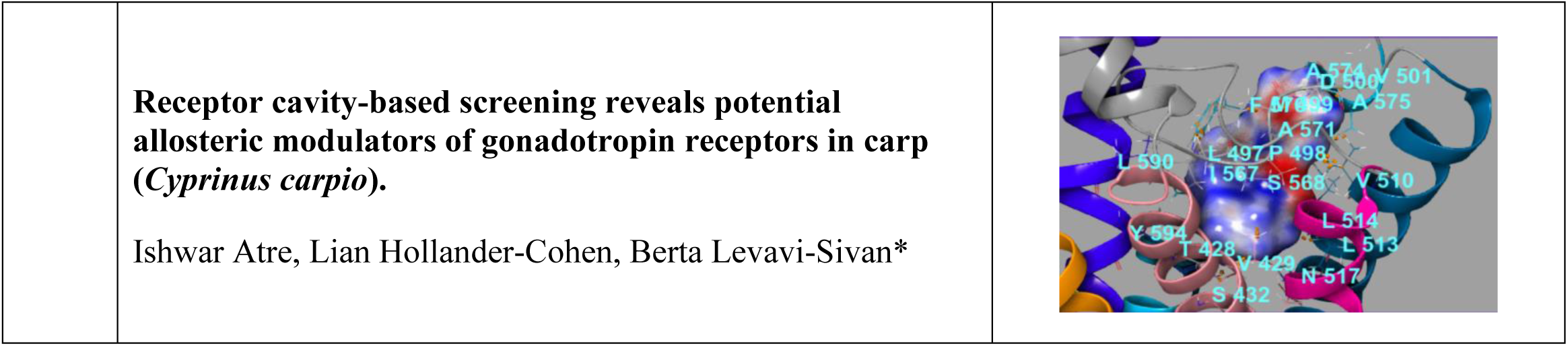

